# Elevated dietary ω-6 polyunsaturated fatty acids induce reversible peripheral nerve dysfunction that exacerbates comorbid pain conditions

**DOI:** 10.1101/2020.05.15.077164

**Authors:** Jacob T. Boyd, Peter M. LoCoco, Ashley R. Furr, Michelle R. Bendele, Meilinn Tram, Qun Li, Fang-Mei Chang, Madeline E. Colley, Dominic A. Arris, Erin E. Locke, Stephan B.H. Bach, Alejandro Tobon, Shivani B. Ruparel, Kenneth M. Hargreaves

## Abstract

Chronic pain is the leading cause of disability worldwide^1^ and commonly associated with comorbid disorders^2^. However, the role of diet in chronic pain is poorly understood. Of particular interest is the Western-style diet, enriched with ω-6 polyunsaturated fatty acids (PUFAs) that accumulate in membrane phospholipids and oxidize into pronociceptive oxylipins^3,4^. Here we report that mice administered a diet enriched with ω-6 PUFAs develop persistent nociceptive hypersensitivities, spontaneously-active and hyper-responsive glabrous afferent fibers, and histologic markers of peripheral nerve damage reminiscent of a peripheral neuropathy. Linoleic and arachidonic acids accumulate in lumbar dorsal root ganglia, with increased liberation via elevated PLA2 activity. Pharmacological and molecular inhibition of PLA2g7 or diet reversal with high ω-3 PUFAs attenuate nociceptive behaviors, neurophysiologic abnormalities, and afferent histopathology induced by high ω-6 intake. In addition, ω-6 accumulation exacerbates the intensity or duration of allodynia observed in preclinical inflammatory and neuropathic pain models, as well as in clinical diabetic neuropathy. Collectively, these data reveal diet as a novel etiology of peripheral neuropathy and risk factor for chronic pain, and implicate multiple therapeutic considerations for clinical pain management.

---

Although medical recommendations about diet are made for cardiovascular disease^5^, diabetes^6^, and autoimmune diseases^7^, this is not the case for most pain disorders. Poor nutrition certainly could be a risk factor for chronic pain conditions, especially with excess intake of ω-6 PUFAs, including linoleic acid (LA) and arachidonic acid (AA). Cellular membrane levels of these essential fatty acids are regulated by dietary intake and necessitate ∼1% of total calories. The average daily Western diet however contains 10-20-fold greater ω-6 levels^8,9^. This discrepancy may have considerable clinical significance since ω-6 PUFAs undergo oxidation into pronociceptive oxylipins^10-14^. Elevated ω-6 levels are associated with pain conditions such as irritable bowel syndrome^15^, rheumatoid arthritis^16^, and headache^17^. Therefore, we evaluated the role of dietary ω-6 PUFAs in the development of persistent pain. We fed mice elevated dietary ω-6 levels, similar to those in the standard American diet, and evaluated changes to baseline nociceptive behaviors as well as those following inflammatory insult or neuropathic injury. We also investigated two therapeutic approaches to counteract diet-induced nociceptive changes.

To determine whether elevated dietary ω-6 PUFAs affects nociceptive thresholds, we administered either a high ω-6 diet (H6D) composed of 11% kcal/kg ω-6 PUFAs or a low ω-6 diet (L6D) composed of 0.75% kcal/kg ω-6 PUFAs to male and female mice for 24 weeks (Supplemental Table 1). Strikingly, both males and females on the H6D developed persistent hypersensitivities to mechanical and heat stimulation, which quickly peaked by 8 weeks (Extended Data Fig. 1). Additional testing at 8 weeks revealed that H6D mice exhibit mechanical hypersensitivity across a range of stimulus intensities, including dynamic brush-evoked stimulation (Fig. 1a,b). H6D mice also demonstrated hypersensitivity to noxious cold in addition to heat (Fig. 1c,d).

**Figure 1.**
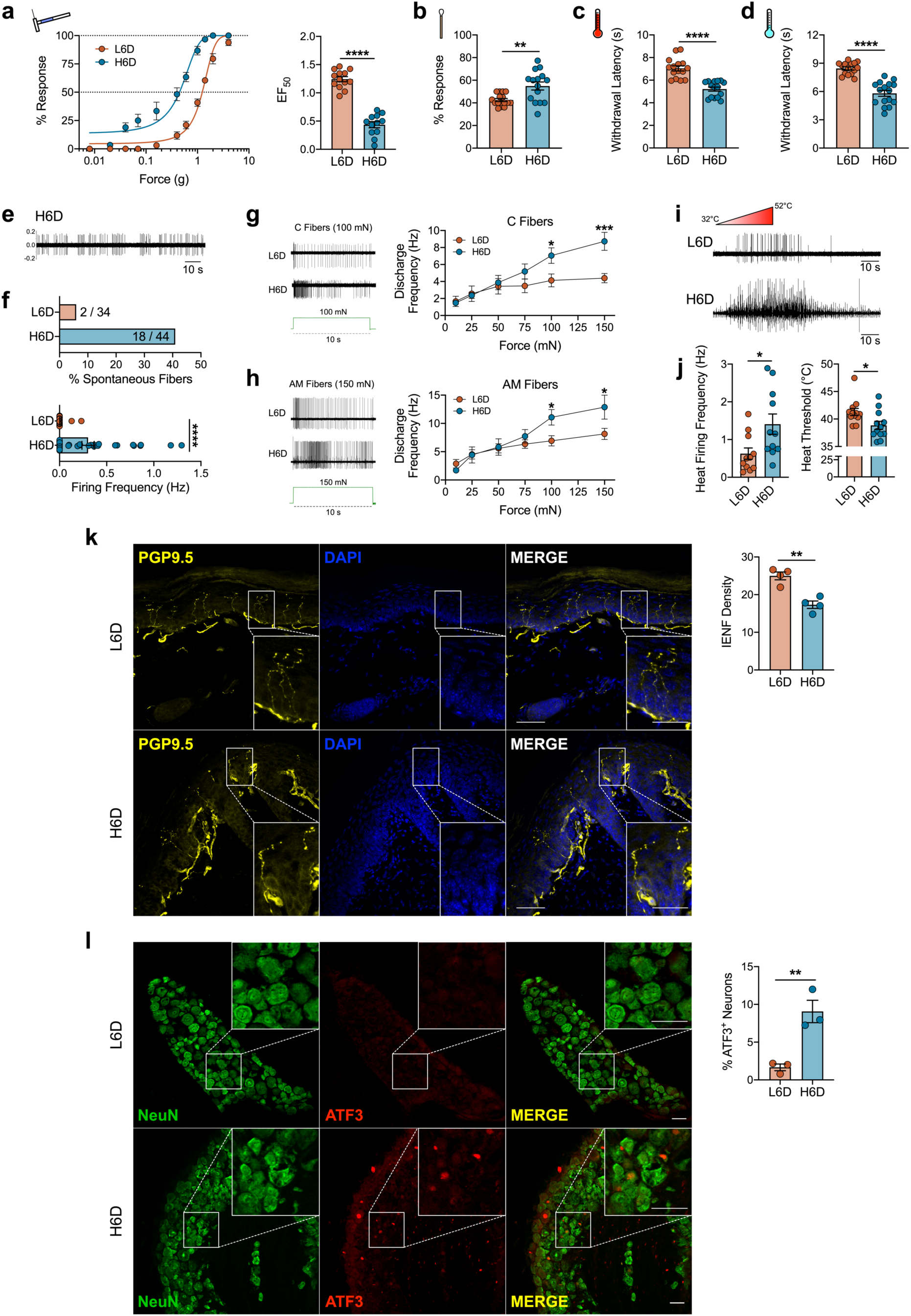
An ω-6 fatty acid-enriched diet induces a peripheral neuropathy-like phenotype in mice. (**a**–**d**) Effects of a high 10% ω-6 fatty acid diet (H6D) or low 0.75% ω-6 fatty acid diet (L6D) on nociceptive thresholds after 8 weeks. (**a**) Mechanical force-response curves (left) and EF_50_ values (right) as determined with nonlinear regression. (**b**) Responsiveness to dynamic brush stimulation. (**c**,**d**) Paw withdrawal latencies (s) to (**c**) radiant heat and (**d**) cold stimulation, n=13-15 mice/group. (**e**) Representative recording of spontaneous firing activity from a H6D mouse. (**f**) Percentage of spontaneously active fibers with mean discharge frequencies (Hz), n=34-44 recordings/group. (**g**,**h**) Representative firing activities of (**g**) C fibers and (**h**) AM fibers responding to 100-mN and 150-mN square force stimulation for 10 s (in green), respectively. Mean discharge frequencies for each force tested to the right for each fiber subtype, n=15-24 fibers. (**i**) Representative firing activities during a heat ramp. (**j**) Heat thresholds and mean discharge frequencies of heat-responsive fibers, n=11-12 fibers/group. (**k**) Representative immunofluorescent staining of PGP9.5 (yellow) with DAPI (blue) to visualize intraepidermal nerve fibers (IENFs) in glabrous hindpaw skin, scale bar: 50 μm. Insets: Designated boxed regions magnified 2.5-fold, scale bar: 25 μm. Mean IENF densities (right), n=4 mice/group. (**l**) Representative immunofluorescent staining of activating transcription factor 3 (ATF3, red) expression in NeuN^+^ lumbar DRG perikarya (green), scale bar: 50 μm. Insets: Designated boxed regions magnified 2.5-fold, scale bar: 25 μm. Mean percent ATF3^+^ neurons (right), n=3 mice/group. ****P<0.0001, ***P<0.001, **P<0.01, *P<0.05 vs L6D. Bars represent group means ± SEM. Each symbol represents data from individual animals or fiber recordings.

We performed single-fiber electrophysiologic recordings from ex vivo glabrous skin-tibial nerve preparations to characterize the effect of the H6D on the detection and firing properties of the terminal endings of peripheral afferent fibers. Interestingly, we found that >40% of fibers from H6D mice exhibit spontaneous firing greater than 0.1 Hz compared to L6D mice (Fig. 1e,f). The H6D increased mechanical-evoked activity in C- and A-fibers as well as post-stimulus afterdischarge (Fig. 1g,h and Extended Data Fig. 2a,b). Moreover, heat-activated fibers from H6D mice had reduced activation thresholds, increased firing frequency, and prolonged post-stimulus activity (Fig. 1i,j and Extended Data Fig. 2c,d). The H6D-induced hyper-responsiveness of afferent fibers to mechanical and heat stimuli parallels the mechanical- and heat-evoked hypersensitivities observed behaviorally in the same plantar hindpaw tissue.

We subsequently evaluated histologic markers of peripheral nerve damage to determine the presence of diet-induced neuronal damage. H6D mice exhibited a significant reduction in intraepidermal nerve fiber (IENF) density, an established marker of preclinical and clinical peripheral neuropathy^18,19^, in glabrous hindpaw skin (Fig. 1k). Further, a significantly higher number of DRG neurons from H6D mice expressed the neuronal stress marker and immediate early gene, activating transcription factor 3 (ATF3)^20,21^ compared to L6D mice (Fig. 1l). Together, the reduction in glabrous IENF density and up-regulation of ATF3 in DRG demonstrate the onset of peripheral nerve damage in mice within 8 weeks on the H6D. In total, the behavioral, neurophysiologic, and pathohistologic data indicate that mice quickly develop a peripheral neuropathy-like phenotype when fed a western-style H6D.

Given that high-fat diet consumption is commonly associated with dyslipidemia, insulin insensitivity, and glucose dysregulation^22-26^, we tested whether the H6D triggered the onset of diabetes and a subsequent neuropathy. Blood glucose as well as HbA1c levels from H6D mice were comparable to mice given the L6D or normal chow, while db/db diabetic mice exhibited elevated levels of both (Extended Data Fig. 3a,b). Additionally, weekly food consumption and body weights in H6D mice were no different than those observed in L6D mice (Extended Data Fig. 3c,d). Therefore, the observed neuropathy-like phenotype does not result from the induction of diabetes.

Rats previously given a H6D showed LA and AA accumulation in multiple tissues, including brain^27^. Since diet-induced nociceptive behaviors can be peripherally- and/or centrally-mediated^13,28-31^, we next evaluated changes to lipid composition for lumbar DRG and spinal cord of H6D mice after 8 weeks using unbiased shotgun lipidomics. Striking changes were observed across lipid classes in lumbar DRG, but not spinal cord, for both male and female mice on H6D compared to L6D mice (Extended Data Fig. 4a,b). We quantified total ω-6 lipid accumulation in DRG and spinal cord, and found that both LA and AA levels were elevated only in the DRG of H6D mice (Fig. 2a,b). Sub-profiling revealed non-uniform accumulation of LA and AA across lipid classes, with the most robust increases observed in membrane-associated lipids (Extended Data Fig. 4c). Interestingly, we also observed marked increases in lysophospholipid levels in DRG of H6D mice (Fig. 2c). Lysophospholipids arise almost exclusively from enzymatic cleavage of sn-2 fatty acids by phospholipase A2 (PLA2) enzymes, and are known to be elevated in diabetes and coronary heart disease,^32^ but also neuropathic pain^33^. The release of sn-2-localized LA and AA from membrane phospholipids by PLA2 initiates their conversion into oxidized, pronociceptive metabolites^34^. Thus, we hypothesized that increased release of ω-6 fatty acids by PLA2 in DRG neurons elevates production of pronociceptive metabolites that underlie the H6D-associated neuropathic phenotype.

**Figure 2.**
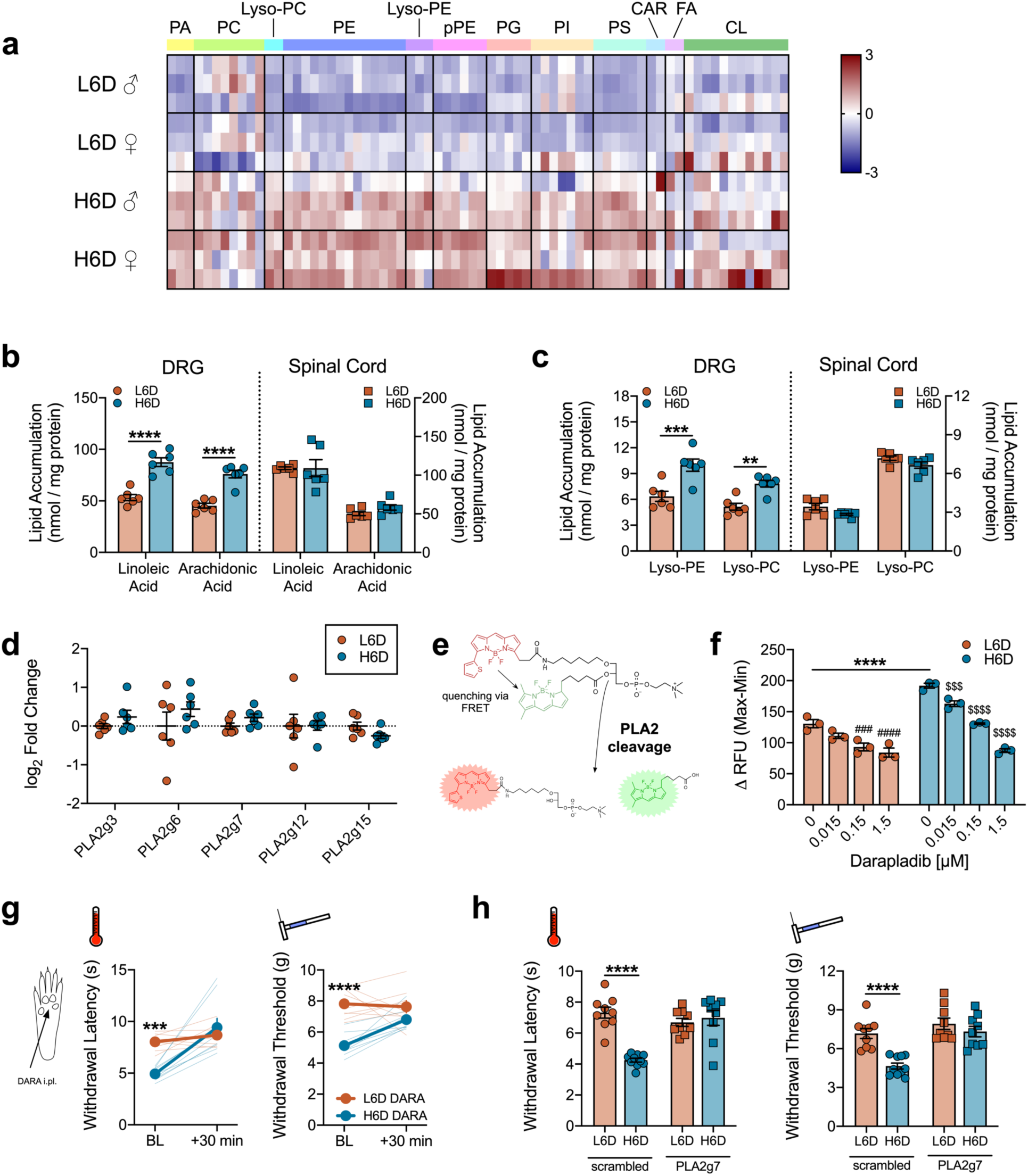
The H6D increases membrane loading of ω-6 PUFAs and stimulates PLA2 activity in peripheral afferent neurons. (**a**) Heatmap of linoleic acid (LA) and arachidonic acid (AA)-esterified lipid species in lumbar DRG from male (♂) and female (♀) mice on either the H6D or L6D, n=3 mice/group. Lipid classes are designated atop the heatmap. Scale bar represents z-score transformations for each lipid species. (**b**,**c**) Total accumulation of (**b**) LA/AA and (**c**) lysophospholipids in lumbar DRG and spinal cord of mice on H6D and L6D. ****P<0.0001, ***P<0.001, **P<0.01 vs L6D, n=6 mice/group. (**d**) PLA2 isozyme expression in lumbar DRG, n=6 mice/group. (**e**) Schematic detailing changes to a fluorogenic BODIPY substrate following sn-2 bond cleavage by PLA2 enzymes. (**f**) PLA2 activity in DRG homogenates in response to pretreatment with vehicle (DMSO) or the PLA2g7-selective inhibitor, darapladib. ****P<0.0001 vs L6D vehicle, ^*####*^P<0.0001, ^*###*^P<0.001 vs L6D vehicle, ^*$$$$*^P<0.0001, ^*$$$*^P<0.001 vs H6D vehicle, n=3 mice/group. (**g**) Acute effects (30 min) of intraplantarly-injected darapladib on heat- and mechanical-evoked nociception. Bold lines represent group means ± SEM determined from individual animal responses (faint lines) before and after treatment, n=8-9 mice/group. (**h**) Withdrawal responses to heat and mechanical stimulation following daily intrathecal (i.t., q.d. × 3 days) injection with either scrambled or PLA2g7-directed siRNA, n=9-10 mice/group. ****P<0.0001, ***P<0.001 vs L6D. Bars represent group means ± SEM. Each symbol represents data from individual animals.

The expression and activity levels of PLA2 isozymes govern the release of membrane lipid-bound LA and AA^35^. We first assessed whether the H6D altered PLA2 isozyme expression in lumbar DRG. Previous single-cell RNA-sequencing identified PLA2g7 as the most prominent isoform in lumbar DRG neurons^36^, accounting for 70-90% of PLA2 transcripts across all sensory neuron subtypes (Extended Data Fig. 5a). We replicated these findings with qPCR using whole DRG RNA extracts as well as by immunolabeling for PLA2g7 expression across several established afferent neuron subclasses (Extended Data Fig. 5b-d). Surprisingly, expression of PLA2g7 and other prominent PLA2 isozymes in DRG were unchanged in both male and female H6D-fed mice (Fig. 2d).

Elevated LA content has been associated with increased PLA2 activity, including in a recent clinical trial where increased PLA2G7 activity was observed in plasma from subjects on a 8-week LA-rich diet^37^. We evaluated for functional alterations in PLA2 activity in afferent neurons from H6D mice by exposing purified DRG homogenates to a phospholipid reporter containing dual fluorogenic boron dipyrromethene (BODIPY) acyl chains (Fig. 2e). Indeed, we found that PLA2 activity was elevated in H6D mice (Extended Data Fig. 6a), thereby suggesting a potential mechanism for increased release of LA and AA in DRG. To assess the contribution of PLA2g7, DRG lysates were pre-incubated with the selective inhibitor, darapladib^38^, before BODIPY exposure. We observed a concentration-dependent decrease in PLA2 activity to an 80% maximal inhibition, revealing that PLA2g7 mediates the majority of PLA2 activity in the DRG (Fig. 2f and Extended Data Fig. 6b,c). Lipidomic analysis determined that ω-6 lipids also accumulate in the glabrous skin of H6D mice, like in the DRG (Extended Data Fig. 6d). Therefore, we next tested whether inhibition of PLA2g7 activity in glabrous skin attenuates H6D-induced nociceptive behaviors. We found that administration of a local, intraplantar injection of darapladib dose-dependently reversed both heat and mechanical hypersensitivity in H6D mice (Fig. 2g and Extended Data Fig. 6e). To validate selective inhibition of PLA2g7 with darapladib, we administered intrathecal siRNA to knock down DRG PLA2g7 in vivo (Extended Data Fig. 6f,g). As with darapladib, PLA2g7 knockdown completely reversed the H6D-induced heat and mechanical hypersensitivities, whereas no effect was seen with scrambled siRNA (Fig. 2h). No effect of either treatment was observed in L6D mice. These data indicate that blocking the first step in oxylipin generation through inhibition of the predominate PLA2 isozyme in lumbar DRG was sufficient to reverse H6D-induced hypersensitivity.

Since blocking PLA2-mediated membrane lipid release reversed the hypersensitivity, we next tested whether balancing lipid membrane content through dietary intervention could reverse the phenotype. It is well-documented that ω-3 PUFA oxylipins exhibit anti-inflammatory and anti-nociceptive effects, which are known to directly counter the pro-inflammatory, pronociceptive effects generated by oxidized metabolites of ω-6 PUFAs^39-41^. It is unclear however, if re-establishing balance between ω-6 and ω-3 membrane lipids with diet could reverse the H6D-induced nerve damage and neuropathy phenotype. To test this, mice were fed the H6D for 8 weeks, at which point they either continued on the H6D or were switched to a high ω-3 diet (H3D) (Fig. 3a and Supplemental Table 1). Remarkably, switching to the H3D completely rescued behavioral heat and mechanical hypersensitivities within 8 weeks (Fig. 3b,c). We next evaluated the effect of altered diet on the electrophysiologic properties of glabrous afferent neuron terminals. H3D mice exhibited a significant, but partial reduction in spontaneously-active fibers compared to H6D mice (Fig. 3d). Interestingly, both C- and AM-fibers from the H3D mice demonstrated recoveries in mechanical responsiveness, heat thresholds, and post-stimulus discharge (Fig. 3e-g and Extended Data Fig. 7a-b). Hindpaw IENF densities and ATF3 expression also recovered to control levels (Fig. 3h,i and Extended Data Fig. 7c,d). Furthermore, LA and AA levels as well as PLA2 activity were reduced in DRG of H3D mice compared to H6D (Fig. 3j,k). Combined, these data demonstrate that a H3D rescues the behavioral, electrophysiologic, pathohistologic, and metabolic alterations associated with the H6D-induced peripheral neuropathy.

**Figure 3.**
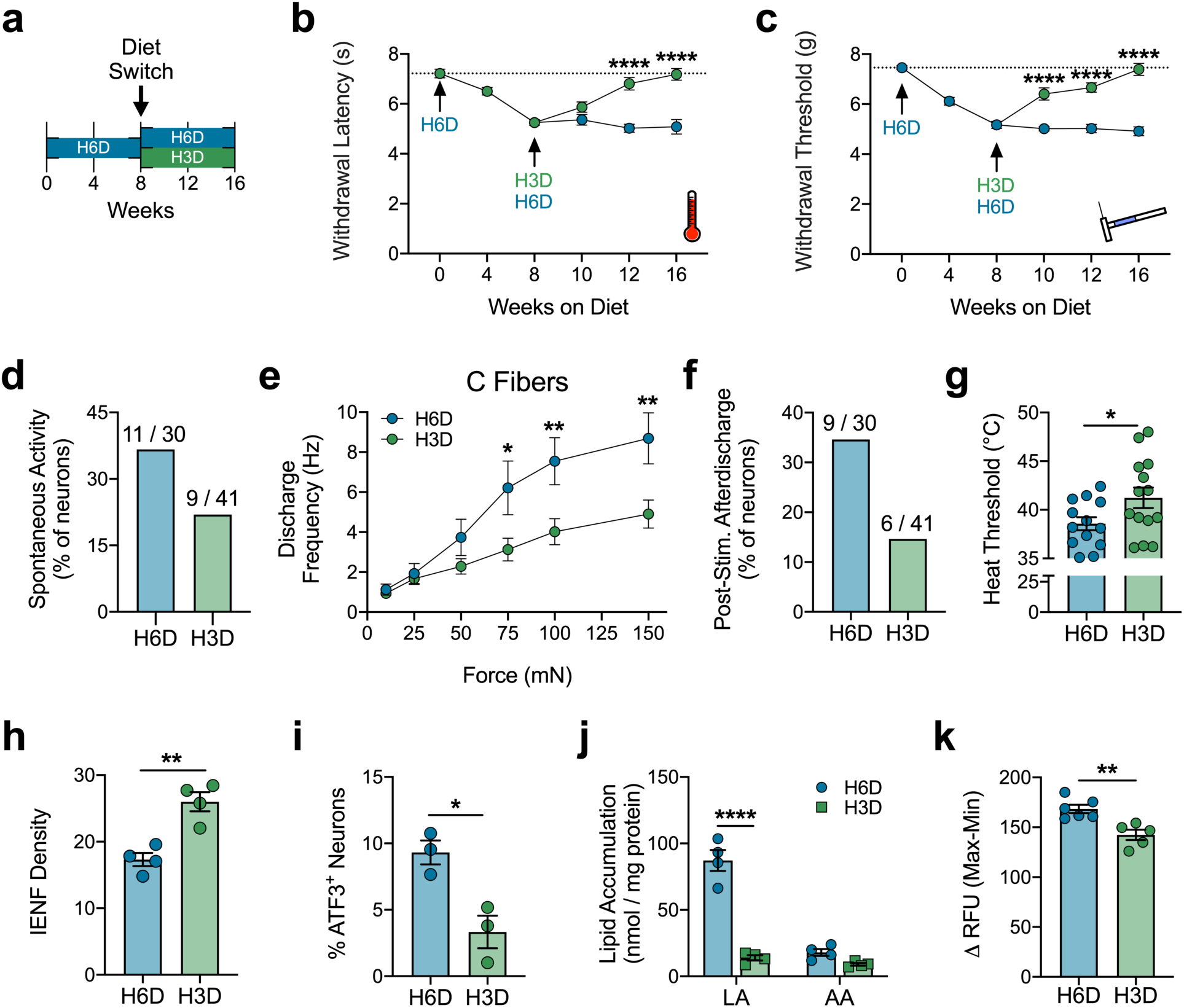
An ω-3 fatty acid-enriched diet rescues the H6D-induced neuropathy-like phenotype. (**a**) Schematic of the diet reversal paradigm, showing mice after 8 weeks on the H6D either continue on the H6D or receive a high 7.3% ω-3 diet (H3D) for 8 additional weeks. (**b**,**c**) Effects of the H3D reversal on H6D-induced (**b**) heat and (**c**) mechanical hypersensitivities, n=11-22 mice/group. (**d**) Percentage of spontaneously-active fibers. Values represent spontaneous fibers over total fibers recorded. (**e**) Mean discharge frequencies of mechanically-responsive C fibers during stimulation, n=14-15 fibers/group. (**f**) Percentage of fibers exhibiting post-stimulus afterdischarge following mechanical force application. (**g**) Heat thresholds of heat-responsive fibers, n=13-14 fibers/group. (**h**) Mean IENF densities, n=4 mice/group. (**i**) Percentage of ATF3^+^ neurons in lumbar DRG, n=3 mice/group. (**j**) Total accumulation of LA and AA in mouse lumbar DRG, n=4 mice/group. (**k**) PLA2 activity in DRG homogenates, n=5-6 mice/group. ****P<0.0001, ***P<0.001, **P<0.01, *P<0.05 vs H6D.

It is well-documented clinically that the development of one pain condition markedly increases the risk to develop additional, more pronounced pain comorbidities^42^. To investigate the effect of diet on hyper-responsiveness to heat and mechanical stimuli, we sought to determine whether the H6D exacerbates nociceptive hypersensitivity and/or prolongs chronicity under inflammatory or neuropathic conditions. To model persistent inflammatory pain^43^, L6D, H6D, and H3D mice were injected with Complete Freund’s Adjuvant (CFA). The H6D prolonged inflammatory heat and mechanical hypersensitivities 3-fold (7 days to 21 days) compared to L6D and H3D mice (Fig. 4a,b). We utilized db/db mice to model type 2 diabetes-associated peripheral neuropathy^44,45^. Diabetic mice on the H6D for 6 weeks developed a greater mechanical allodynia compared to mice on the L6D, whereas db/db mice on the H3D did not develop mechanical allodynia at all (Fig. 4c). Interestingly, db/db mice given either the L6D or H3D exhibited normal responses to noxious heat stimulation, contrasting the hypersensitivity observed with the H6D (Fig. 4d). Together, these results indicate that a H6D is a risk factor for prolonged and/or exacerbated nociception associated with an inflammatory or neuropathic injury, but can be ameliorated with a H3D.

**Figure 4.**
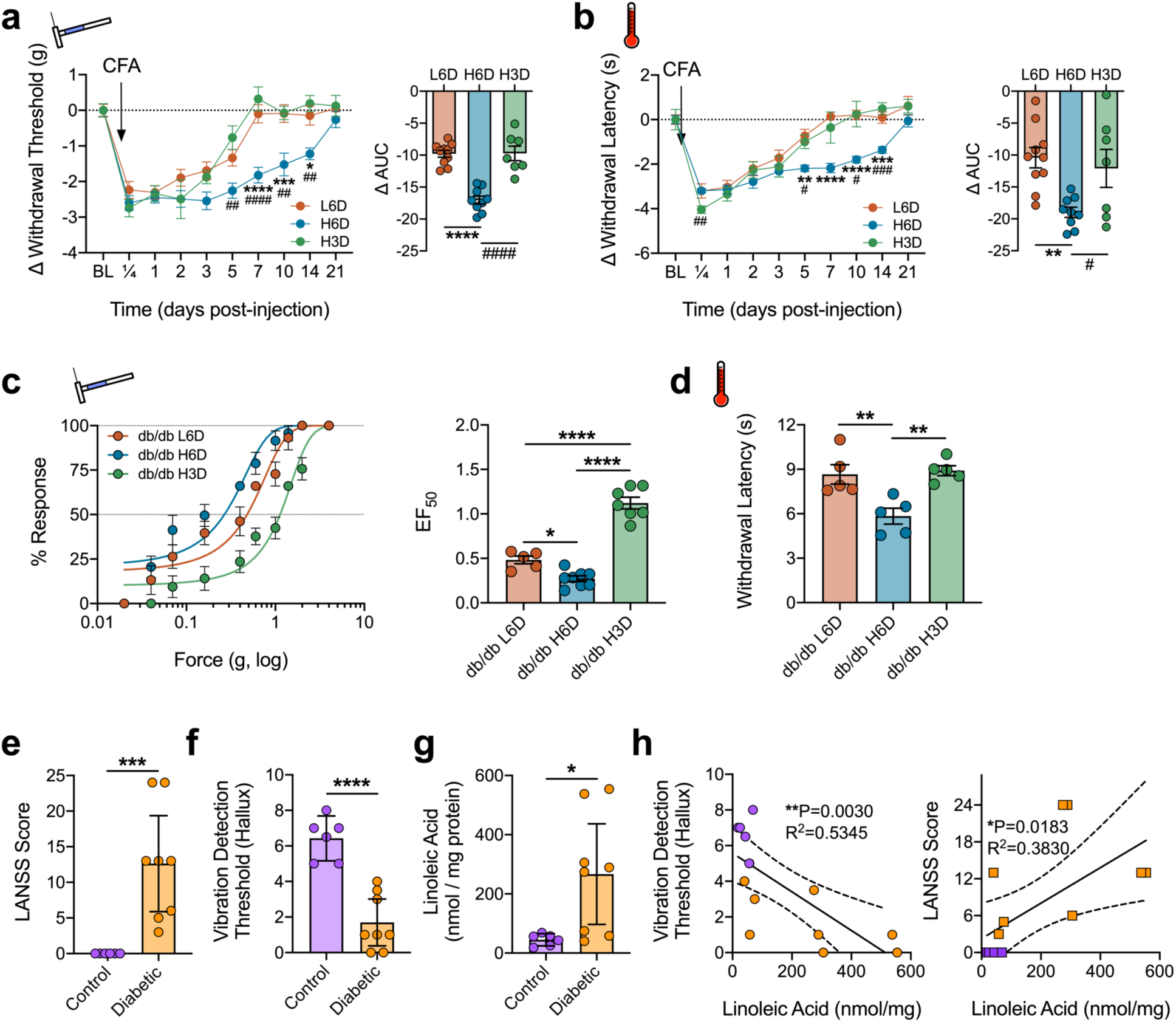
Diet-specific modulation of nociceptive behaviors associated with inflammatory and neuropathic pain. (**a**,**b**) Time course of L6D-, H6D-, and H3D-specific changes to (**a**) mechanical paw withdrawal thresholds (g) and (**b**) heat paw withdrawal latencies (s) following administration of Complete Freund’s Adjuvant (CFA). Change from baseline (BL) was calculated for each mouse since mean pre-CFA BLs differed across groups. Insets: Areas under the curve (AUC). ****P<0.0001, ***P<0.001, **P<0.01, *P<0.05 L6D vs H6D, ^####^P<0.0001, ^###^P<0.001, ^##^P<0.01, ^#^P<0.05 H6D vs H3D, n=7-10 mice/group. (**c**,**d**) Mechanical force-response curves and EF_50_ values for 12-week old db/db mice fed either a L6D, H6D, or H3D for 6 weeks. (**d**) Paw withdrawal latencies (s) to radiant heat stimulation. ****P<0.0001, **P<0.01, *P<0.05, n=5 mice/group. (**e**,**f**) Scatter plots of (**e**) LANSS Pain Scale scores and (**f**) vibration detection thresholds (a.u.) of the right hallux for control (purple) and diabetic neuropathy (orange) patients. (**g**) Total LA content in skin biopsies from the lateral malleolus. Horizontal bars represent group means ± 95% confidence intervals. ****P<0.0001, **P<0.01, *P<0.05 vs control, two-tailed Mann-Whitney test for (**e**), two-tailed Student’s t test with Welch’s correction for (**f**,**g**), n=6-8 patients/group. (**h**) Correlation analysis between patient skin LA levels and their respective hallux vibration detection thresholds and LANSS scores (Pearson, two-tailed, ***P=0.001).

Since these findings could have far-reaching clinical significance, we tested for an association between skin ω-6 PUFA levels and pain symptoms in patients with diabetic neuropathy. Based on the LANSS Pain Scale^46^, diabetic neuropathy patients exhibited a significant increase in neuropathic pain-related symptoms, including allodynia, compared to age-matched controls (Fig. 4e). Diabetic patients also showed a marked reduction in vibration detection thresholds of the halluces (Fig. 4f). Lipidomic assessment revealed that LA content was elevated in ankle skin biopsies of diabetic patients (Fig. 4g and Extended Data Fig. 8), and, despite the small sample size, robust, significant correlations between LA levels and vibration detection thresholds (R^2^=0.5345) and LANSS scores (R^2^=0.3830) were observed (Fig. 4h).

Our findings support the inclusion of pain disorders with other prominent diseases that necessitate medical oversight of patients’ diet. We demonstrate that the typical Western diet loads DRG and skin with ω-6 PUFAs within just 8 weeks, leading to elevated PLA2-mediated lipid release and the onset of peripheral nerve damage and nociceptive hypersensitivity. We also propose diet as a pain risk factor, based on the observations that elevated ω-6 PUFAs exacerbate nociceptive hypersensitivities in preclinical models of inflammatory and neuropathic pain as well as in patients with diabetic neuropathy. These are certainly consistent with high-BMI patients being increasingly susceptible to chronic pain conditions^47,48^. We further show that administration of darapladib, a PLA2G7-selective inhibitor under clinical investigation for non-pain conditions^49,50^, attenuates H6D-induced hypersensitivity, while dietary replacement with ω-3 PUFAs completely reverses the H6D phenotype. Together, these may provide novel approaches for treating or preventing chronic pain disorders.

## Acknowledgments

These studies were supported by the National Institutes of Health grants R01 NS110948 (K.M.H.), UL1 TR002645 (K.M.H), T32 DE14318 (P.M.L., A.R.F.), T32 GM113896 (J.T.B.), F30 AT009949 (J.T.B.), F32 DK113841 (P.M.L.), F30 DE028486 (A.R.F.), and a grant from the Ella and Williams Owen’s Foundation (K.M.H.). Certain mass spectrometric analyses were carried out on equipment supported by the U.S. Department of Agriculture, Agricultural Research Service, under agreement No. 58-3094-8-012. This manuscript is solely the responsibility of the authors and does not necessarily represent the official views of the NIH or U.S. Department of Agriculture. We thank Xianlin Han and lab for expertise and guidance on the shotgun lipidomics. We thank Mayur Patil and Ping Wu for technical assistance as well as Anibal Diogenes, Nikita Ruparel, Asma Khan, and Armen Akopian for fruitful discussions.

## Author contributions

J.T.B., P.M.L., S.R., and K.M.H. conceived and designed the studies; J.T.B., P.M.L, M.B., and M.T. conducted the behavioral experiments; A.R.F., P.M.L., and F.C. conducted the single-fiber electrophysiology; J.T.B. and P.M.L performed the histology; Q.L., F.C., D.A.A., and P.M.L. performed the BODIPY experiments; P.M.L. D.A.A., and M.T. performed total tissue lipid extractions; M.E.C. and S.A.B. conducted the quantitative LC-MS/MS; P.M.L. and F.C. performed western blots; E.L. and A.T. conducted neurological assessments on trial participants and collected skin punch biopsies; J.T.B., P.M.L., and K.M.H. performed data analysis for all experiments, J.T.B. and P.M.L. prepared the figures, images, and illustrations; J.T.B., P.M.L., and K.M.H. wrote the manuscript; all authors revised the manuscript.

## Competing financial interests

The authors declare no competing financial interests.

## Materials and Methods

### Animals

All animal experiments conformed to the Guidelines for the Use of Animals in Research as put forward by the International Association for the Study of Pain, and to protocols approved by the University Texas Health Science Center at San Antonio (UTHSCSA) Animal Care and Use Committee. Mouse experiments were initiated at 8-10-weeks of age in both male and female C57BL/6J mice (#000664, The Jackson Laboratory). Male BKS.Cg-Dock7^m^+/+Lepr^db^/J (db/db^51^, #000642, The Jackson Laboratory) mice 16-weeks of age were used as positive controls to model a diabetic phenotype.

### Diets

Randomized groups of mice were fed isocaloric diets containing 10% g/kg total fat with an energy density of 22.9% kcal/kg for at least 8 weeks to maintain equivalency of nutrient and calorie intake. L6D, H6D, and H3D were formulated by Dyets Inc. (Bethlehem, PA) based on the AIN-93G diet^52^, and modified to control the amount of ω-6 or ω-3 PUFAs (Supplementary Table 1). As described previously^27^, the H6D consisted of 11.5% kcal/kg ω-6 PUFAs, 0.9% kcal/kg ω-3 PUFAs, 2.4% kcal/kg MUFAs, and 7.4% kcal/kg SFA (Dyet #181189) and the L6D consisted of 0.3% kcal/kg ω-6 PUFAs, 0.9% kcal/kg ω-3 PUFAs, 2.3% kcal/kg MUFAs, and 18.8% kcal/kg SFA (Dyet #180784). H3D consisted of 0.5% kcal/kg ω-6 PUFAs, 7.3% kcal/kg ω-3 PUFAs, 7.0% kcal/kg MUFAs, and 6.1% kcal/kg SFA (Dyet #104593). For rescue experiments, mice were placed on H6D for 8 weeks, then switched to either H6D or H3D for at least another 8 weeks. All diets were stored at −20°C and used for no longer than 6 months. Food was replaced weekly. Animals and food were weighed weekly to monitor changes in body weights and food consumption. Weekly food intake (per cage of 5 mice) was calculated by the change in food weight measured at the beginning and end of each week. Normal chow consisting of 2.6% kcal/kg ω-6 PUFAs, 0.3% kcal/kg ω-3 PUFAs, 1.3% kcal/kg MUFAs, and 0.8% kcal/kg SFA (Teklad LM-485; Envigo) was given to control and db/db mice for metabolic comparison.

### Behavioral testing

All experiments were performed by blinded observers and the assay order was randomized on each testing day.

#### Mechanical stimulation assays

Paw withdrawal thresholds (PWT) to noxious mechanical stimulation were evaluated using Ugo Basile dynamic plantar aesthesiometer equipped with an 0.8-mm rigid von Frey filament as previously described^53^. Briefly, animals were randomized into plastic observation boxes on an acrylic grid platform and acclimated for 60 min. The aesthesiometer was positioned under the mouse to stimulate the mid-plantar area of the hindpaw with a force ramp up to 15 g over a 7 sec period. The force at which withdrawal occurred was recorded. All animals were tested a minimum of three times and the average was used for statistical analysis. In order to more thoroughly characterize mechanical hypersensitivity, a range of von Frey fibers from 0.008–4 g (11 fibers) were used as described previously^54^. Briefly, mice were acclimated in observation boxes on a wire mesh floor for 60 min. Von Frey fibers contacted the midplantar surface of the hindpaw until slight buckling of the filament was observed, then held for 2 sec. Each fiber was probed 5 times per mouse with at least 30 sec between applications. Withdrawal of the paw was noted as a positive response and no movement was a negative. Values were recorded as percent response for each fiber. Force-response curves were generated using nonlinear least-squares regression of the mean and used to calculate the half maximal effective force (EF_50_)^55^.

#### Brush test

Dynamic tactile response was assessed using a cotton swab brushed quickly across the plantar surface of the hindpaw as described previously^56^. Animals were acclimated for 30-60 minutes in boxes atop mesh grid flooring. The “puffed out” cotton swab was used to brush the length of the ventral hindpaw in a continuous motion. Positive responses were defined as withdrawal or rapid shaking of the paw. Each animal was tested 5 times on each paw with a minimum of 30 sec between tests.

#### Heat stimulation assay

Paw withdrawal latency (sec) to heat stimulation was assessed using the radiant heat test^57^. Animals were acclimated in plastic observation boxes for 30-60 min prior to testing. The mid-plantar surface of the mouse hindpaw was exposed to a radiant heat source through a glass floor until paw withdrawal. The intensity of the heat source was adjusted to produce a standard baseline of ∼8 sec in naïve wild type mice, with a maximum cutoff of 20 sec. All animals were tested a minimum of three times with at least 60 sec between recordings and the average of all recordings was used for statistical analysis. Mechanical thresholds were determined from the same animals on the same days.

#### Cold stimulation assay

Response to cold stimulus was measured according to an adaptation of a previously described protocol^58^. Mice were placed in plastic observation boxes on top of a 3/16” tempered glass flooring and allowed to acclimate for at least 30 min. A 10-mL syringe was sectioned above the Leur-lock and tightly packed with finely crushed dry ice. The syringe was pressed firmly on the bottom of the tempered glass directly below the hindpaw while measuring paw withdrawal latency (sec) with a stop-watch. Thickness of tempered glass and syringe size were selected in order to establish baseline measurements of ∼8 sec. Each animal was tested 3 times on each paw with a minimum of 1 min between tests.

#### Complete Freund’s Adjuvant (CFA) pain model

Complete Freund’s adjuvant (CFA; Sigma; St. Louis, MO) was diluted 1:1 with saline and injected intraplantarly (i.pl.) into the right hindpaw of mice using a 30G insulin syringe filled to 20 µl. Thermal and mechanical readings were taken prior to the injection and then at 0.25, 1, 2, 3, 5, 7, 10, 14, and 21 days after injection. Animals were followed until they returned to the original baseline thresholds. Since pre-CFA baselines were different between diet cohorts, data were normalized as the change from baseline withdrawal response.

### Blood sampling

HbA1c and fasting blood glucose levels were measured in mice after 8 weeks on either H6D, L6D, or H3D. Whole blood was collected from the submandibular branch of the jugular vein into EDTA-coated Microvette CB300 collection tube (Kent Scientific). HbA1c levels were determined using a mouse whole blood assay kit according to the manufacturer’s instructions (#83010, Crystal Chem). Briefly, whole blood was mixed with lysis buffer for 10 minutes then mixed with protease buffers and incubated at 37°C for 5 minutes. Absorbance was then measured at 700-nm on a microplate reader (Molecular Devices; San Jose, Ca, VersaMax). To measure fasting blood glucose, mice first were fasted for 5 hours to optimize physiological context^59,60^. Then, a single drop of blood was collected from the submandibular branch of the jugular vein on a blood glucose test strip and analyzed with the AlphaTrak2 Blood Glucose Monitoring System (Parsippany, NJ). Blood from WT and db/db mice on normal chow served as a negative and positive controls, respectively.

### Shotgun lipidomics

DRG and spinal cord tissues were homogenized in 0.5 mL 10x diluted PBS in 2.0 ml cryogenic vials (Corning Life Sciences) by using the Precellys® Evolution (Bertin Corp). Protein assay on the homogenates was performed by using a bicinchoninic acid protein assay kit (Thermo Scientific) with bovine serum albumin as standards. The rest of the homogenate was accurately transferred into a disposable glass culture test tube, and a mixture of lipid internal standards was added prior to lipid extraction for quantification of all reported lipid species. Lipid extraction was performed by using a modified Bligh and Dyer procedure as described previously^61^. Individual lipid extracts were resuspended into a volume of 400 μl of chloroform/methanol (1:1, v/v) per mg of protein and flushed with nitrogen, capped, and stored at −20°C for lipid analysis. For shotgun lipidomics, lipid extracts were further diluted to a final concentration of ∼500 fmol/µL, and the mass spectrometric analysis was performed on a QQQ mass spectrometer (Thermo TSQ Quantiva) equipped with an automated nanospray device (TriVersa NanoMate, Advion Bioscience) as previously described^62^. Identification and quantification of lipid molecular species were performed using an automated software program^63,64^. Data were normalized to protein (per mg). All membrane-bound lipids (defined as lipids attached to membrane-anchoring headgroups) were then grouped together and the total concentration of the ω-6 fatty acids, linoleic acid (18:2) and arachidonic acid (20:4), were determined between L6D and H6D and used for statistical analysis.

### Quantification of total tissue lipids

The lipid extraction protocol was adapted from the Bligh-Dyer method^65^. Briefly, frozen tissue samples were weighed, then sectioned on a cryostat at 16 µm and collected into a cold-acclimated glass test tube. Two ml extraction buffer (50% methanol, 25% chloroform, 20% ddH2O, 0.02% butylated hydroxytoluene) was added to each tube on ice, spiked with internal standards (5 µl/ml d_4_-LA, 1 µl/ml d_8_-AA, 5 µl/ml d_6_-ALA, 1 µl/ml d_5_-EPA, 1 µl/ml d_5_-DHA), and vortexed for 30 sec every 5 min for 15 min. Samples were centrifuged at 5000 × g at 4°C for 10 min, and the bottom organic phase was collected in a separate glass tube. The aqueous phase was re-extracted with 1 ml extraction buffer, and following collection of the second bottom phase, samples were dried down under a steady stream of nitrogen. To analyze total lipid pools, samples next underwent base-catalyzed saponification as described previously^66,67^. Briefly, dried samples were re-suspended with 850 µl methanol/chloroform solution (8:1) and 150 µl 40% potassium hydroxide solution, placed under nitrogen, then heated to 37°C for 60 min. Following, samples received 700 µl 0.05M phosphate buffer and 300 µl 2.5M HCl (pH <5) then were extracted twice with 2 ml hexane. Both upper phases were combined then dried down under nitrogen. Samples were stored at −80°C until processing with LC-MS/MS.

Dried samples were re-constituted with 150 µl ethanol and transferred to 1 ml autosampler vials for LC-MS/MS. A Waters Acquity UPLC system was used to perform reversed-phase separation of the free fatty acids. A Thermo Electron Corporation (Bellefonte, PA) BDS Hypersil C18 column (50 mm × 2.1 mm i.d.) with a 5-μm particle size was held at 45°C. Solvent A was 5 mM ammonium acetate in water and solvent B was acetonitrile with 10% 2-propanol and 0.2% acetic acid. All solvents were Fisher Scientific Optima LC/MS grade (Fair Lawn, NJ). The gradient was set-up as follows: 0 min – 60% A:40% B, 5 min – 60% A:40% B, 8 min – 5% A: 95% B, 9 min – 5% A; 95% B, 9.10 – 80% A:20% B, 11 min – 80% A:20% B. The autosampler was held at 5 °C. The injection volume was 5 μl and the flow rate was constant 0.5 mL/min. A TQD tandem quadrupole mass spectrometer (Waters Corporation, Milford, MA) was utilized for free fatty acid concentration determination. An ESI source in negative ion mode was used with the capillary voltage set to 3 kV. The source temperature was set at 120°C with a desolvation temperature of 250°C. Argon was used as the collision gas for CID at 0.10 mL/min and nitrogen was used for the cone gas flow at 1 L/hr and the desolvation gas flow at 500 l/hr. Multi-reaction monitoring (MRM) channels were used for both LA and AA, but since LA does not produce detectable fragment ions, the channel utilized the parent ion mass of 279.2 for the fragment ion as well. This will exclude species of the same mass that do produce fragment ions. AA MRM transitions were 303.2→205, 303.2→259, 303.2→285. The collision energy for LA was 1 eV and for AA was 20 eV. Waters Corporation software MassLynx was used to perform channel integration and smoothing. Analytical standards of AA and LA were ordered from Cayman Chemical (Ann Arbor, MI) and used to establish calibration curves. The calibration ranged from 10 ppm to 10 ppb for each standard in ethanol.

### Single-fiber recordings from glabrous skin-tibial nerve preparations

To provide a neurophysiological correlate to H6D-induced behaviors, we utilized ex vivo glabrous skin-tibial/sural nerve preparations as described previously^68-72^. Tibial and sural nerves were selected based on anatomical distribution of terminal nerve endings within the glabrous skin of the mouse hindpaw^73^, using the same region tested for evoked behavioral assays described above. Mice were briefly anesthetized with isofluorane and then sacrificed via cervical dislocation. The right leg was shaved followed by careful dissection of the skin-nerve preparation, which included the glabrous skin and attached tibial/sural nerves. The preparation was transferred to the organ bath chamber of a 3D-printed stage (courtesy of Peter Reeh) with oxygenated standard interstitial fluid (SIF, pH 7.4) consisting of: 123 mM NaCl, 3.5 mM KCl, 2.0 mM CaCl_2_, 0.7 mM MgSO_4_, 1.7 mM NaH_2_PO_4_, 9.5 mM NaC_6_H_11_O_7_, 5.5 mM glucose, 7.5 mM sucrose, 10 mM HEPES, and perfusing at 32 ± 0.7°C (Eco Silver, LAUDA-Brinkmann) at a flow rate of 15-16 ml/min (Masterflex L/S, Cole-Parmer). The skin was positioned corium-side up and pinned down with insect needles to a silicon rubber base (Sylgard 184, Dow Corning) in the organ chamber. The nerve was threaded through a 1-mm hole into the adjacent recording chamber and placed atop a mirror plate. Low viscosity mineral oil (M5904, Sigma) was applied onto the nerve on the mirror plate during teasing and recording to electrically isolate it from the rest of the perfusion chamber. Using a stereomicroscope (SMZ 745T, Nikon) positioned over the recording chamber, the nerve was de-sheathed and single filaments were carefully teased apart for recording. Teased filaments were wrapped around a 0.25 mm silver wire electrode (AGW1010, World Precision Instruments) to record activity. Both the reference (grounded extracellular milieu) and recording (nerve) electrodes in the recording chamber fed into a low-noise headstage probe (DAM80p, World Precision Instruments) which relayed the signal to a DAM80 differential amplifier (World Precision Instruments). Incoming signals were amplified 1000-fold and bandpass-filtered between 300 and 1000 Hz. Amplified signals were filtered through a Hum Bug 50/60 Hz Noise Eliminator (Quest Scientific) then passed to a TBS1000B digital oscilloscope (Tektronix), a Model 330 audio monitor (A-M Systems), and a Micro1401-3 digital data acquisition system (Cambridge Electronic Design). Data were recorded with Spike2 (Cambridge Electronic Design).

#### Sensory profiling of teased fibers

Receptive fields (RF) were initially identified based on mechanical responsiveness to a blunt glass rod. Upon positive identification of a RF, the following steps were utilized to characterize the sensory profile of the teased fiber: (1) *spontaneous activity –* The presence of spontaneous activity was assessed for the first 2 min of recording, (2) *mechanical stimulation –* A precision force-controlled mechanical stimulator (Series 300C-I Dual Mode Servo System, Aurora Scientific; Aurora, Ontario) was used to evaluate fiber mechano-sensitivity. The mechanical cylinder (0.7 mm tip diameter) was positioned over the RF of interest, followed by computer-controlled application of square force stimuli (force range of 5 – 200 mN with 10 sec duration). To prevent sensitization/desensitization of the recorded fiber, 60 sec intervals were given between force applications, (3) *heat stimulation –* A Peltier-based thermal stimulus delivery system (CS1, Cool Solutions) was used to assess fiber responsiveness to controlled heat application^74^. A custom-designed cylindrical ring that was 3D-printed on a Form 2 using Tough Resin (FormLabs) was used to isolate the RF of interest. Vacuum grease (Dow Corning) tightly sealed the ring in position. An insulated afferent tube connected to the Peltier device, an efferent tube, and a thermocouple were then positioned within the ring. A dispensing pump (IPC24, Ismatec) drove the push-pull superfusion (60 ul/sec) of SIF over the RF within the ring (Figure 2.2). The SIF temperature increased from 30°C to 52.0-60.5°C during heat ramps (30 sec). After 30 sec of heating, the Peltier was stopped and temperature returned to baseline within 90 sec. At least 5 min intervals were given between heating to prevent sensitization/desensitization of the isolated RF, (4) *conduction velocity –* A 2.0 MΩ Parylene-coated tungsten metal stimulating electrode (TM33B20, World Precision Instruments) was positioned within the RF of interest. A stimulus isolator (A365, World Precision Instruments) and pulse generator (A310, World Precision Instruments) were used to deliver electrical pulses to the RF. Upon electrical stimulation of the fiber, digital calipers were used to measure the conduction distance (in mm) between the RF and the recording electrode. Conduction velocity (CV) was calculated by dividing this distance by the latency of the firing fiber from the stimulating artefact and represented as meters per second (m/s).

#### Spike detection and analysis

Recordings were analyzed with off-line template-matching in Spike2 version 8.14 (Cambridge Electronic Design). Settings for spike template matching required a minimum 70% of sampling points within a template and 25% maximum amplitude change. Waveform data was interpolated linearly. High-pass filter time constant was set to 6.4 ms. Triggers were set at least 3 times the baseline noise level. Conduction velocity (CV), calculated by dividing the conduction distance over the spike electrical latency, was used to classify fiber type according to the following cut-offs: C fibers <0.8 m/s; AM fibers >1.2 m/s^75^. Spontaneously active fibers were defined with a minimum unprovoked discharge frequency of 0.1 Hz^76^. Mechanical discharge frequencies were determined for each force application as the number of firings within the 10 sec ramp period. Post-stimulus after-discharge was identified as persistent activity occurring after completion of any force ramp. For heat responses, total afferent activity was used instead of discharge frequency as recordings often included individual units with variable amplitudes that could not always be discriminated^69^. Heat threshold temperatures were identified as the temperature that initiated the first afferent activity during a heat ramp. Post-stimulation activity was calculated as the number of action potentials during the time period defined as the point when the recovering temperature reaches a fiber’s heat threshold through to baseline.

### Immunohistochemistry and imaging

L3-L5 dorsal root ganglia (DRG) and 3 mm biopsies of glabrous hindpaw skin were immediately dissected after sacrifice. Tissues were immersion-fixed in 4% paraformaldehyde in 0.1M phosphate buffer (PB) for 2 hr. Tissue samples were washed 3 × 15 min in PB, immersed in 10% sucrose at 4°C overnight, transferred to 30% sucrose at 4°C overnight and then stored at −20°C. For cryo-sectioning, tissues were thawed and placed in OCT (Tissue-Tek, Sakura Finetek; Torrance, CA) prior to freezing on dry ice. Sections of DRG (14 μm) and paw tissue (20 μm) were cut with a cryostat (Microm HM550, Thermo Fisher Scientific; Waltham, MA) and thaw-mounted onto Superfrost Plus slides (Fisherbrand; Waltham, MA). Slides were air-dried at RT and stored at −20°C. Staining was performed using MAXpack Immunostaining Media Kit (Active Motif; Carlsbad, CA). Tissue sections were blocked with MAXblock Blocking Medium for 1 h at room temperature, followed by incubation with primary antibodies diluted in MAXbind Staining Medium at 4°C overnight. Primary antibodies included: PGP9.5 (1:1000; AB1761-1, Millipore), ATF3 (1:400; ab207434, Abcam), NeuN (1:500; ABN91, Millipore), NFH (1:5000; 822601, Biolegend), TRPV1 (1:700; GP14100, Neuromics), PLA2g7 (1:500; 15526-1-AP, ProteinTech), IB4 (1:800; I21411, ThermoFisher), GFRA2 (1:500; AF429, R&D Systems). Sections were then washed 3 × 10 min in MAXwash solution and incubated with secondary antibodies for 1 hr at RT. The following secondary antibodies (Jackson ImmunoResearch) were used: Alexa Fluor® 488 AffiniPure Donkey Anti-Chicken IgY (IgG) (H+L), Alexa Fluor® 488 AffiniPure Donkey Anti-Rabbit IgG (H+L), Alexa Fluor® 568 Donkey anti-Rabbit IgG (H+L), Alexa Fluor® 647 AffiniPure Donkey Anti-Guinea Pig IgG (H+L), Rhodamine Red-X AffiniPure Donkey Anti-Goat IgG (H+L) all at 1:500 dilution from stock. After washing 3 × 10 min with MAXwash, DAPI (0.02 ug/ml, Sigma; St. Louis, MO, D9542) was applied for 10 min at RT, followed by 2 × 10 min washes with MAXwash and then 2 × 5 min in ddH_2_O. Slides air-dried in the dark for 20 min then were mounted with ProLong Diamond (ThermoFisher Scientific; Waltham, MA) and no. 1.5 high precision cover slips (Zeiss; Oberkochen, Germany) for imaging.

#### Confocal Microscopy

Images were obtained with a Nikon C1si laser scanning confocal microscope equipped with: 402-nm diode, 488.1-nm solid state, 561.4-nm diode-pumped solid state, and 639-nm diode lasers. Objectives (Nikon; Melville, NY) used include: a 20X 0.75 NA Plan Apo DIC air and 40X 0.95 NA collar-corrected Plan Apo air. Image acquisition settings were 1024 × 1024 resolution, 12-bit image depth, 2.0 μs/pixel scan speed, 30 μm confocal aperture (pinhole diameter), and sequential channel scan. Z-stacks were taken at 0.3-0.6 μm optical steps. All images for each tissue type were taken at identical gain settings. Laser power, HV, and offset were adjusted to maximize dynamic range while avoiding pixel saturation. Adjustments of brightness/contrast, look-up tables, and z-stack reconstructions were performed in FIJI^77-80^.

#### Intraepidermal Nerve Fiber (IENF) Density Quantification

To evaluate changes in epidermal innervation density, immunolabeled IENFs projected across the dermal-epidermal junction to the epidermis were counted in randomly selected fields of view from each section. The following exclusion criteria were used when counting IENFs: (1) fragments of nerve fibers in the epidermis that did not clearly cross the dermal-epidermal junction were not counted and (2) fibers that approached but did not cross the dermal-epidermal junction, as determined by basal epidermal autofluorescence, were not counted^81,82^. To reduce bias, counting was conducted by 2 blinded observers on 3-5 non-consecutive sections per animal with 4 animals per group. The averaged fiber count for each section was divided by the length of the dermal-epidermal junction to calculate the IENF density (IENFs/mm of tissue). Mean IENF density ± SEM for each group was used to determine statistical significance.

#### ATF3 Quantification

To measure changes in the expression of the transcription factor, ATF3, as a marker of neuronal injury^20^, the number of ATF3^+^ neurons in immunolabeled sections from L3-L5 DRG were counted. Images underwent thresholding before manual counting of the stained nuclei. For each image, the percentage of ATF3-labeled neurons was calculated by dividing total ATF3^+^ neurons by total NeuN^+^ neurons × 100. Using FIJI, quantifications were performed on 3-5 non-overlapping sections per mouse to determine individual means. Group means ± SEM were calculated using individual means from 3-4 mice per group.

### PLA2 activity assay

DRGs from H6D, L6D, and H3D animals were dissected, placed into cold lysis buffer (50mM Tris HCl, pH 8, 2mM DTT, 1mM MgCl_2_, complete Mini protease inhibitor) and incubated on ice for 15 minutes. DRGs were homogenized using a 2-ml Dounce homogenizer, transferred to 1.5 ml tubes and centrifuged for 20 min at 15,000g and 4°C. Supernatants were collected into new 2-ml tubes. Total protein concentration was determined using the Bio-Rad Bradford protein assay kit according to the manufacturer’s instructions (ThermoFisher Scientific, #23225). An adapted BODIPY (Invitrogen, #A10072) assay was used for continuous monitoring of phospholipase A2 (PLA2) activity from homogenates based on the manufacturer’s instructions and previous publication. Briefly, 1mM stock of Red/Green BODIPY PC-A2 was prepared in DMSO. To prepare the liposome mixture, equal volumes of 10mM DOPC, 10mM DOPG and 1mM BODIPY dye were mixed in a microcentrifuge tube and slowly injected into assay buffer while rapidly stirring. DRG lysates were added to microplate wells in appropriate concentrations. A buffer-only well was use as a negative control and bee venom (PLA2g3) was used as a positive control. The liposome mixture was then added 1:1 to the microplate wells. In PLA2 inhibitor experiments, the drug was incubated with the DRG lysate for 15 minutes prior to addition of the liposomal substrate. FlexStation 3 (Molecular Devices; San Jose, CA) was set to 37°C and used for kinetic recording of 520 nm and 570 nm emission for 30 minutes. Peak maximum minus minimum fluorescent signals for each wavelength (a.u.) were used for statistical analysis for each sample.

### In vivo treatment

All drugs were prepared in coded syringes on the day of injection by an individual not performing behavioral testing in order to blind the experimenter. Animals were initially acclimated to observation boxes atop thermal or mechanical platforms for a minimum of 30 min prior to collection of baseline recordings. After baselines, animals were administered vehicle or testing compound. Darapladib (Selleckchem; SB-480848, Houston, TX) was dissolved in dimethyl sulfoxide (DMSO) and Tween-20 then diluted in phosphate buffered saline (PBS) to 2% of each for final concentrations of 300 nmol and 30 nmol. Control animals received equivalent volume of the darapladib vehicle.

Knockdown of PLA2g7 was performed using Dharmacon SMARTpool small interfering RNA (siRNA) against four separate PLA2g7 target sequences. Control groups received non-targeting scrambled (scr) siRNA (Dharmacon RNA Technologies, Lafayette, CO). The PLA2g7 or scrambled siRNA was injected intrathecally (i.t., 2 µg) for three consecutive days for a cumulative dose of 6 µg. Behavior was tested on 2 days after final injection by blinded observers, followed by collection of DRGs for protein and transcript quantification.

### qPCR

L3-5 DRG were harvested and immediately frozen on dry ice then stored at −80°C. DRG tissue was homogenized as described previously^83^ in 900 μl QIAzol lysis reagent (QIAGEN; Hilden Germany) and placed on ice for 5 min. RNA was isolated according to the manufacturer’s instructions (Qiagen; RNeasy Plus Universal Mini Kit, #73404). Briefly, 100 µl gDNA eliminator and 180 µl chloroform were added to each tube and shaken for 15 s. Tubes were spun at 12,000 g for 15 min and the top aqueous phase was removed for further purification with ethanol and wash buffers and elution with RNase-free water. cDNA was then synthesized from 1 μg RNA using Superscript III First Strand Synthesis kit (Invitrogen; #18080051, San Diego, CA). Amplification of target sequences was performed using TaqMan Fast Advanced Master Mix (Applied Biosystems; #4444557, Foster City, CA) according to the manufacturer’s instructions plus selected primers for Rn18S (Mm03928990_g1, ThermoFisher), PLA2g3 (Mm00555594_m1, ThermoFisher), PLA2g6 (Mm00479527_m1, ThermoFisher), PLA2g7 (Mm00479105_m1, ThermoFisher), PLA2g12 (Mm01316982_m1, ThermoFisher), and PLA2g15 (Mm00505425_m1, ThermoFisher). qPCR was performed using the StepOnePlus Real-time PCR System (Applied Biosystems, #4376600). A no-template control (NTC) was used as a negative control. Cycles of thresholds (Ct) were collected and normalized to the internal control, Rn18S. Normalized ΔΔCt to Rn18S reference were used for statistical analysis^84^.

### Western Blot

Reinforced screw cap microtubes (2-ml) pre-filled with 2.8 mm ceramic beads (19-628, Omni International) were filled with 900 μl RIPA buffer (ThermoFisher, #89900) with cOmplete mini protease inhibitors (Roche; #11836145001, Basel, Switzerland). Thawed DRG tissue samples were then transferred to bead tubes and placed on ice. The processing chamber of a Bead Ruptor 24 (Omni International; #19-040E, Kennesaw, GA) was pre-cooled to 0°C with an attached BR-Cryo Cooling Unit (Omni International; #19-8005). Samples were placed in the tube carriage, homogenized for 30 sec at 7.10 m/s, then placed on ice for 5 min. Samples were transferred to fresh tubes and centrifuged for 3 min at 8,000 × g and 4°C. Supernatants were collected and stored at −80°C until use. After total protein determination, supernatants were prepared for SDS-PAGE on 4–12% gradient Bis-Tris gels according to the NuPAGE protocol (Novex; San Diego, CA). Proteins were transferred to PVDF membranes via iBlot2 (ThermoFisher). Immunoblots were probed with polyclonal rabbit anti-PLA2G7 antibody (1:500, 15526-1-AP, ProteinTech). Actin was measured as a loading control with a monoclonal mouse anti-actin antibody (1:1000, sc-47778, Santa Cruz; Santa Cruz, CA). Donkey anti-rabbit IR800 and donkey anti-mouse IR680 secondary antibodies (1:10,000, LI-COR Biosciences) were used to detect PLA2G7 and actin, respectively. Immunoblots were imaged using a LI-COR Odyssey infrared imager and relative band intensities were quantified using Image Studio (LI-COR Biosciences).

### Human study and sample collection

Clinical sensory data and skin biopsies collected from 6 control subjects and 8 subjects with type 2 diabetes and diabetic neuropathy (all ages >55 years) were included as part of an ongoing observational trial at the University of Texas Health Science Center at San Antonio. Regulatory approval was obtained from the Human Subjects Institutional Review Board at the University of Texas Health Science Center at San Antonio (#20160027HU). Subject demographics are available in Supplementary Table 2. Informed consent was obtained from subjects according to protocol guidelines set by the Institutional Review Board. Subjects completed the self-reported pain questionnaire, the Leeds Assessment of Neuropathic Symptoms and Signs (LANSS), followed by sensory testing as described^46^. Vibration detection thresholds then were determined for both halluces for each patient, followed by collection of a skin punch biopsy.

For vibration testing, a Rydel-Seiffer tuning fork (US Neurological) was used on the left and right hallux (i.e., great toe). The principal neurologist placed the base of the tuning fork, with the dampers facing the researcher, on the bony prominences. To create a vibration, the tynes were pressed together between the thumb and index finger, then released such that the illusion of two triangles is visible on each damper marked on a scale from 0 to 8. As the intensity of the vibration diminishes, the two triangles move closer together and the point of intersection moves slowly upward. Each patient vocalizes once the vibration cannot be detected and the intensity score was recorded. This procedure was repeated four times per hallux, with the first reading being discarded and the last three being recorded.

Lastly, following injection with 2% lidocaine/epinephrine (Hospira) solution to anesthetize the skin, punch biopsies were taken 10 cm proximal to the lateral malleolus of the ankle to a depth of 4 mm with a sterile 4-mm circular punch. Biopsies were collected in sterile 2-ml cryotubes and immediately snap-frozen on liquid nitrogen. Samples were stored at −80°C until processing for total lipid analysis.

### Data and statistical analysis

GraphPad Prism 8.0 (GraphPad, La Jolla, CA) was used for statistical analysis. Differences between groups were assessed by unpaired Student’s *t*-test, one-way or two-way ANOVA with Tukey’s post-hoc test for pairwise comparison. Statistical significance was established based on a two-sided alpha of 0.05 for all tests. For rigor and transparency, we used general linear model (GLM) methods for analysis (e.g., ANOVA with Bonferroni’s test) or non-linear ANOVA (von Frey test) using sample sizes designed to generate an 80% power at a two-sided P<0.05. All experiments that utilized male and females were tested for possible sex differences and if none were observed, the animals were combined for further analysis^85^. Observers were blinded to treatment allocation, and all manufacturing information of the diets and all tested compounds were retained.

---

**Extended Data Figure 1:**
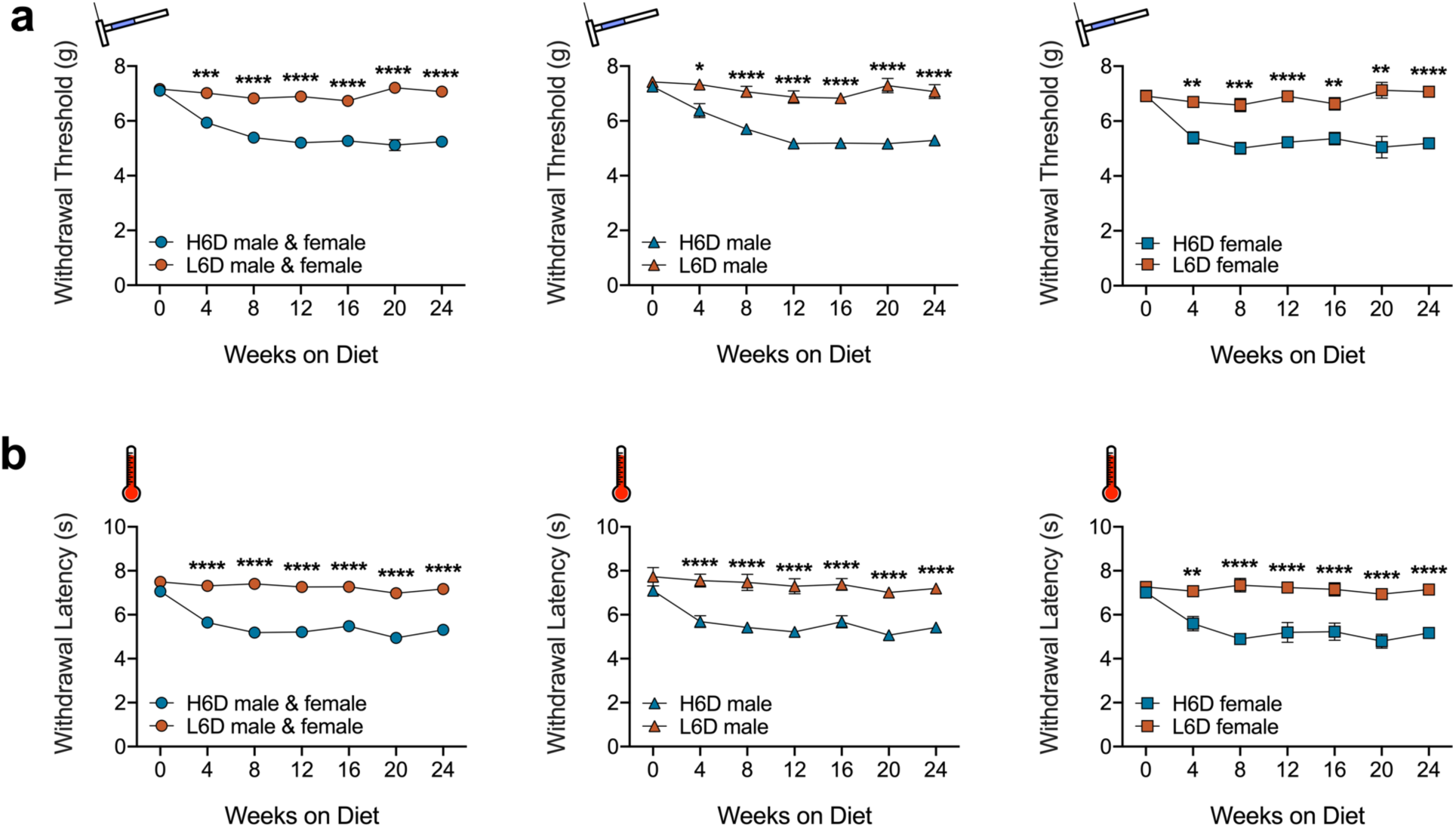
H6D induces persistent nociceptive hypersensitivities in both male and female mice. (**a**) Time course of changes in mechanical withdrawal threshold (g) for mice on H6D and L6D. (**b**) Time course of changes in heat withdrawal latency (s) for mice on H6D and L6D. Left plots are male and female responses compiled together, middle plots are male-only responses, right plots are female-only responses. ****P<0.0001, ***P<0.001, **P<0.01, *P<0.05 vs L6D, n=13-15 mice/group.

**Extended Data Figure 2:**
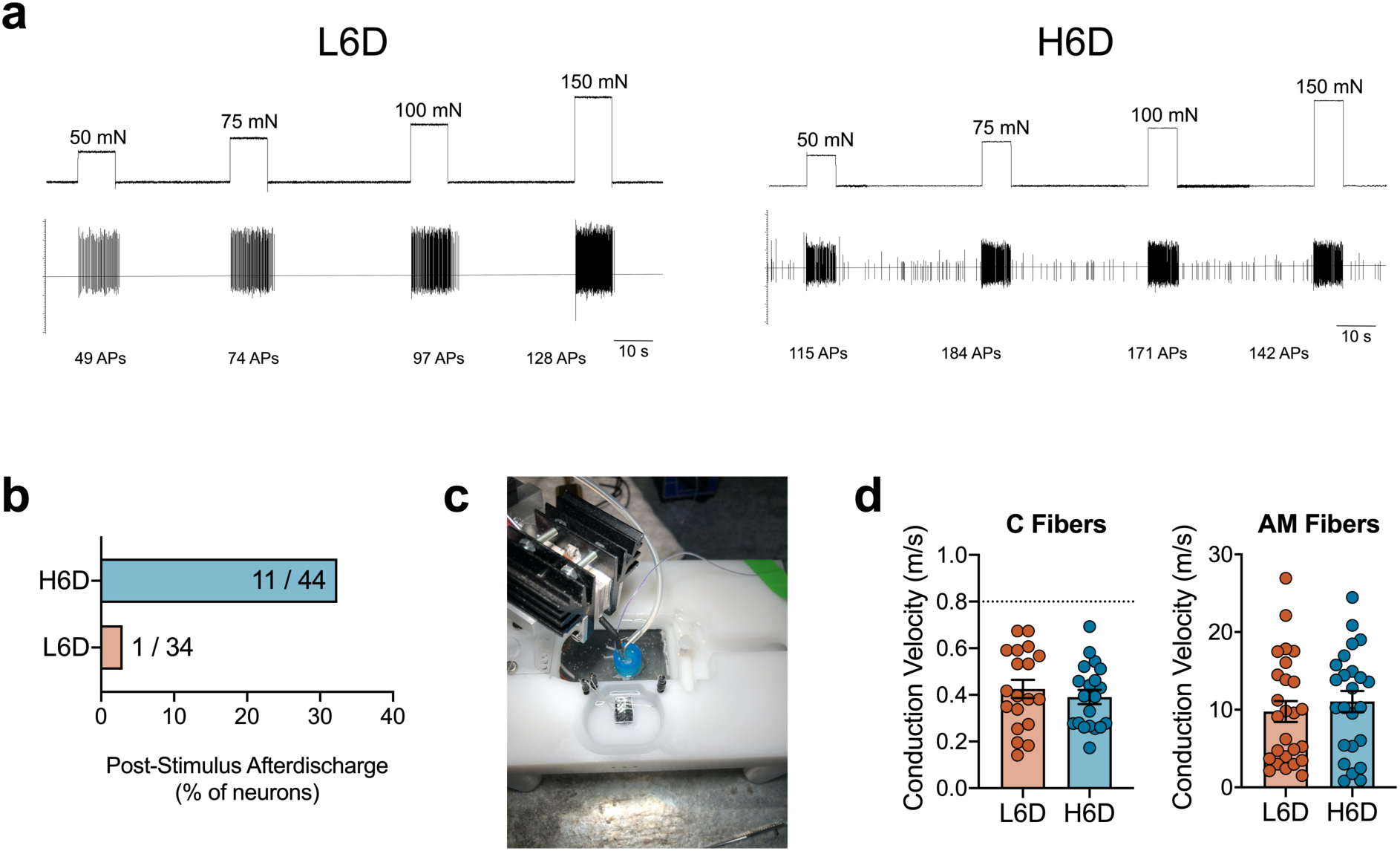
The H6D sensitizes afferent fibers to mechanical and heat stimuli. (**a**) Representative recording wavemarks from L6D and H6D mice during mechanical force application. The number of action potentials are denoted beneath each stimulation for each recording. (**b**) Percentage of fibers exhibiting post-stimulus afterdischarge following mechanical force application. Values represent the number of fibers exhibiting afterdischarge over the total recorded fibers for each group. (**c**) The peltier-based heat delivery system setup. (**d**) Conduction velocities (m/s) determined for recorded C and AM fibers from L6D and H6D mice, n=19-24 fibers/group.

**Extended Data Figure 3:**
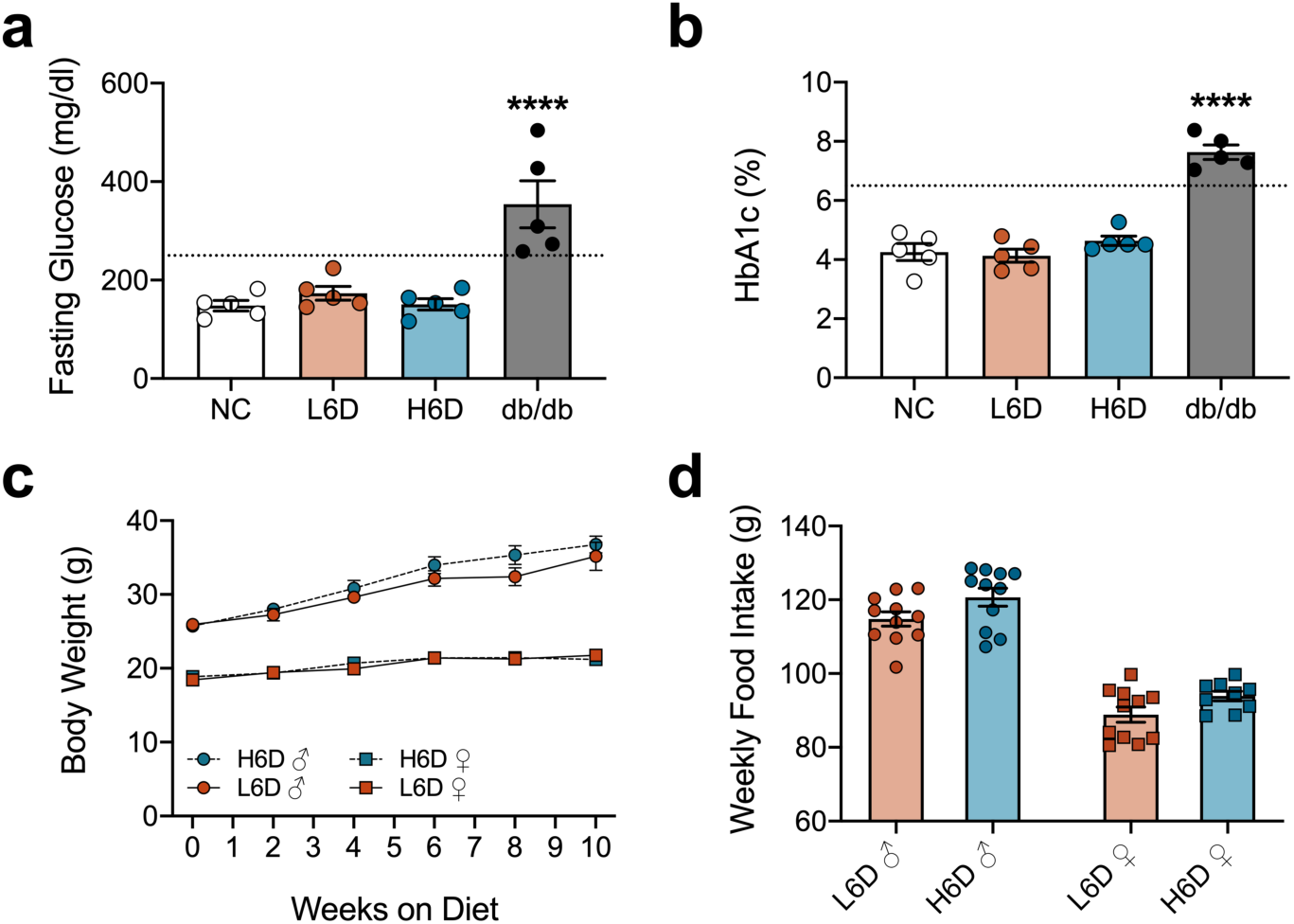
The H6D does not induce a diabetic phenotype. (**a**,**b**) Scatter plots of (**a**) fasting blood glucose levels and (**b**) HbA1c levels from mice on L6D and H6D for 8 weeks. Mice on normal chow (NC) and 16-week-old db/db mice served as the negative control and positive control, respectively. Dotted lines in each figure represent established cut-offs for diabetes. ****P<0.0001 vs NC, n=5/group. (**c**,**d**) Weekly monitoring of (**c**) body weights and (**d**) food intake for both male and female mice on either L6D or H6D.

**Extended Data Figure 4:**
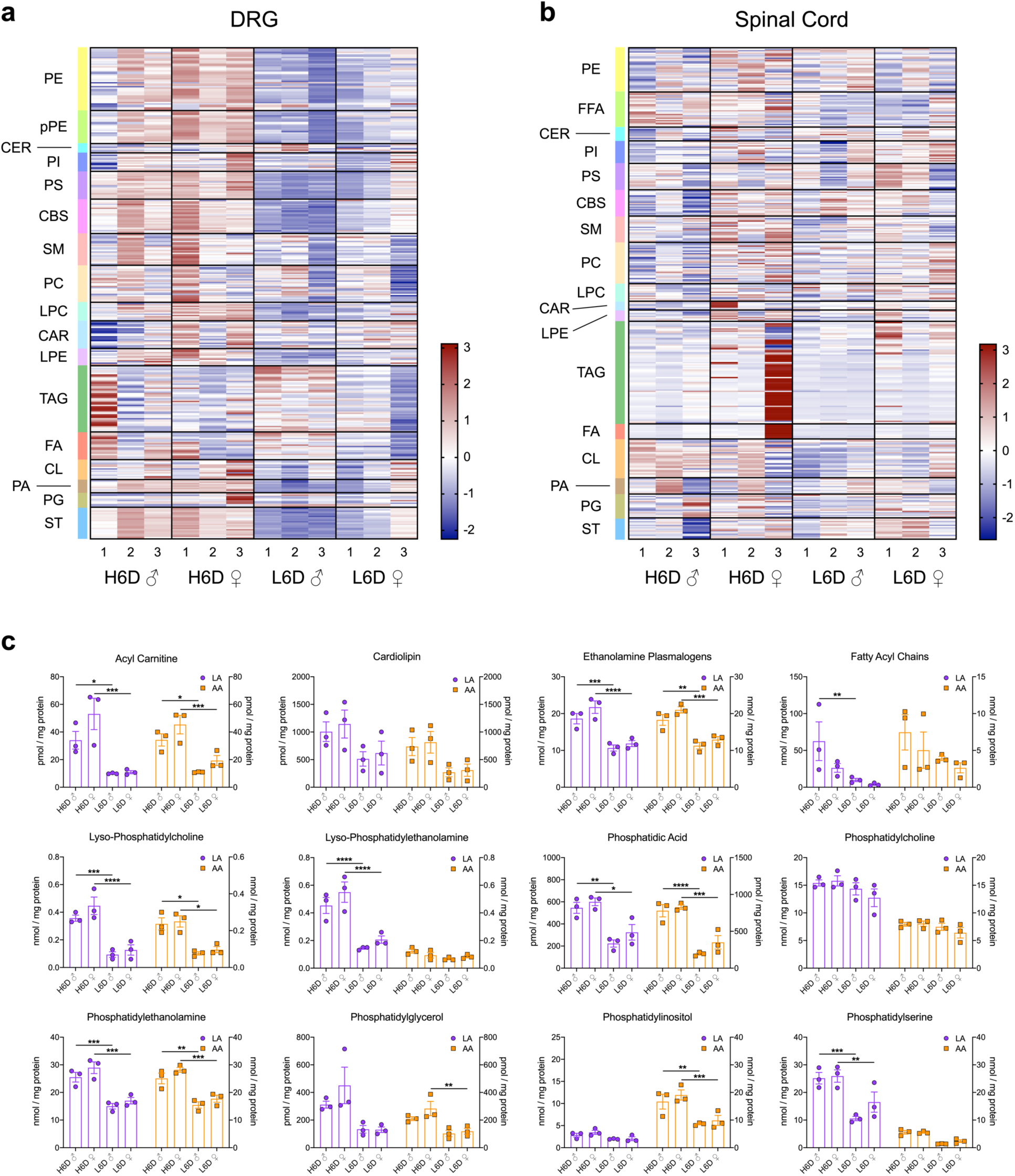
The H6D alters lipid composition in DRG, but not spinal cord. (**a**,**b**) Heatmaps of lipid species from (**a**) lumbar DRG and (**b**) spinal cord from male (♂) and female (♀) mice on either the H6D or L6D. Lipid classes are designated to the left of each heatmap. Scale bar represents z-score transformations for each lipid species. (**c**) LA- and AA-esterified lipids in DRG sub-profiled by lipid class for male and female mice on either the H6D or L6D. ****P<0.0001, ***P<0.001, **P<0.01, *P<0.05 vs L6D (male or female), n=3/group.

**Extended Data Figure 5:**
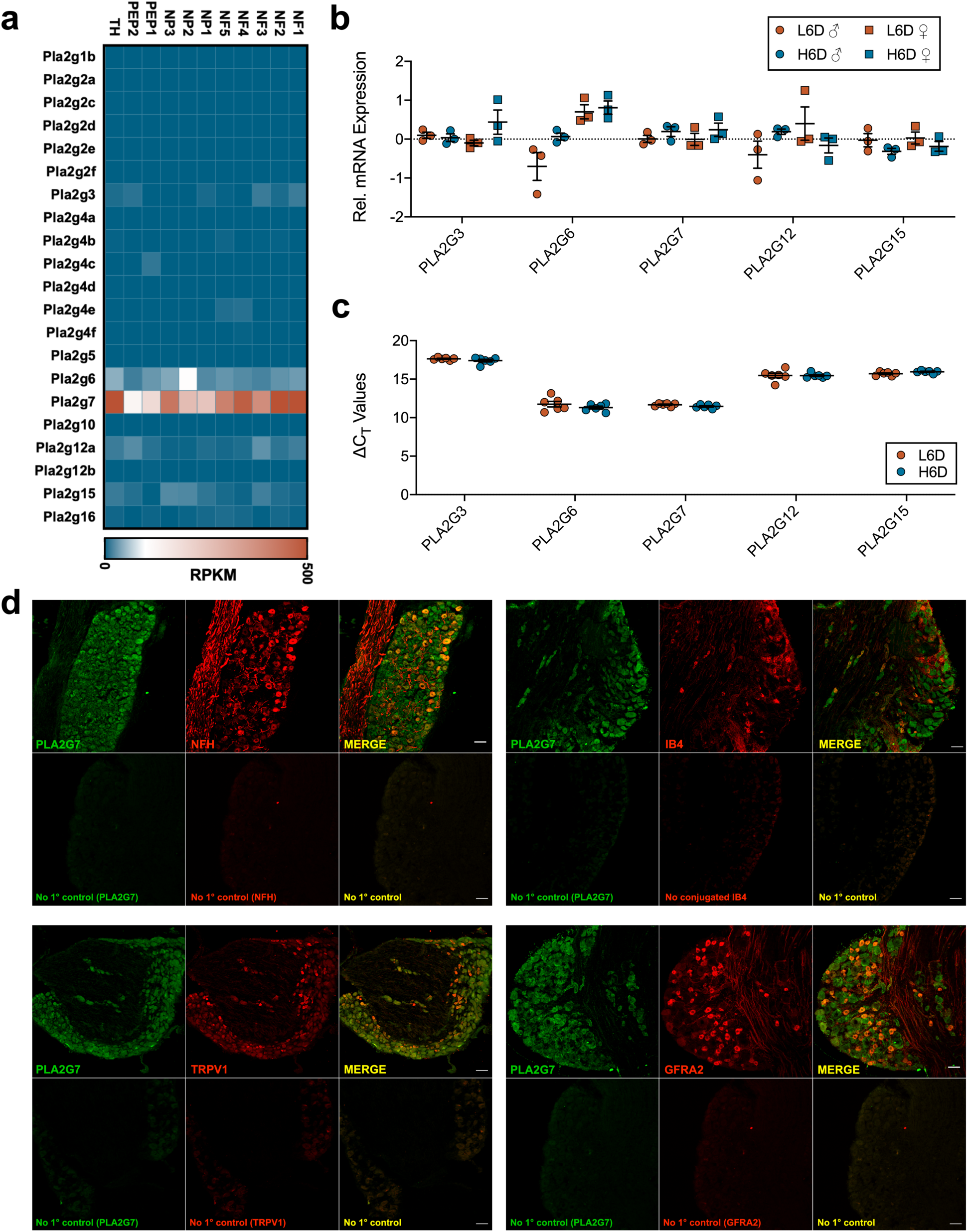
PLA2g7 expression predominates in neuronal subpopulations of the lumbar DRG. (**a**) Heatmap indicating PLA2 isozyme expression across established neuronal subpopulations of the mouse lumbar DRG. Single cell RNA-seq data were reproduced with permission from Usoskin et al., 2015. (**b**,**c**) qPCR data showing (**b**) PLA2 isozyme expression (male and female breakdown) and (**c**) change in cycle threshold values relative to 18S rRNA in lumbar DRG from H6D and L6D mice. (**d**) Representative immunofluorescent staining of PLA2g7 expression in mouse lumbar DRG and co-localization with neuronal subtype-specific markers. No primary controls are included for each marker as designated. Scale bar: 50 μm.

**Extended Data Figure 6:**
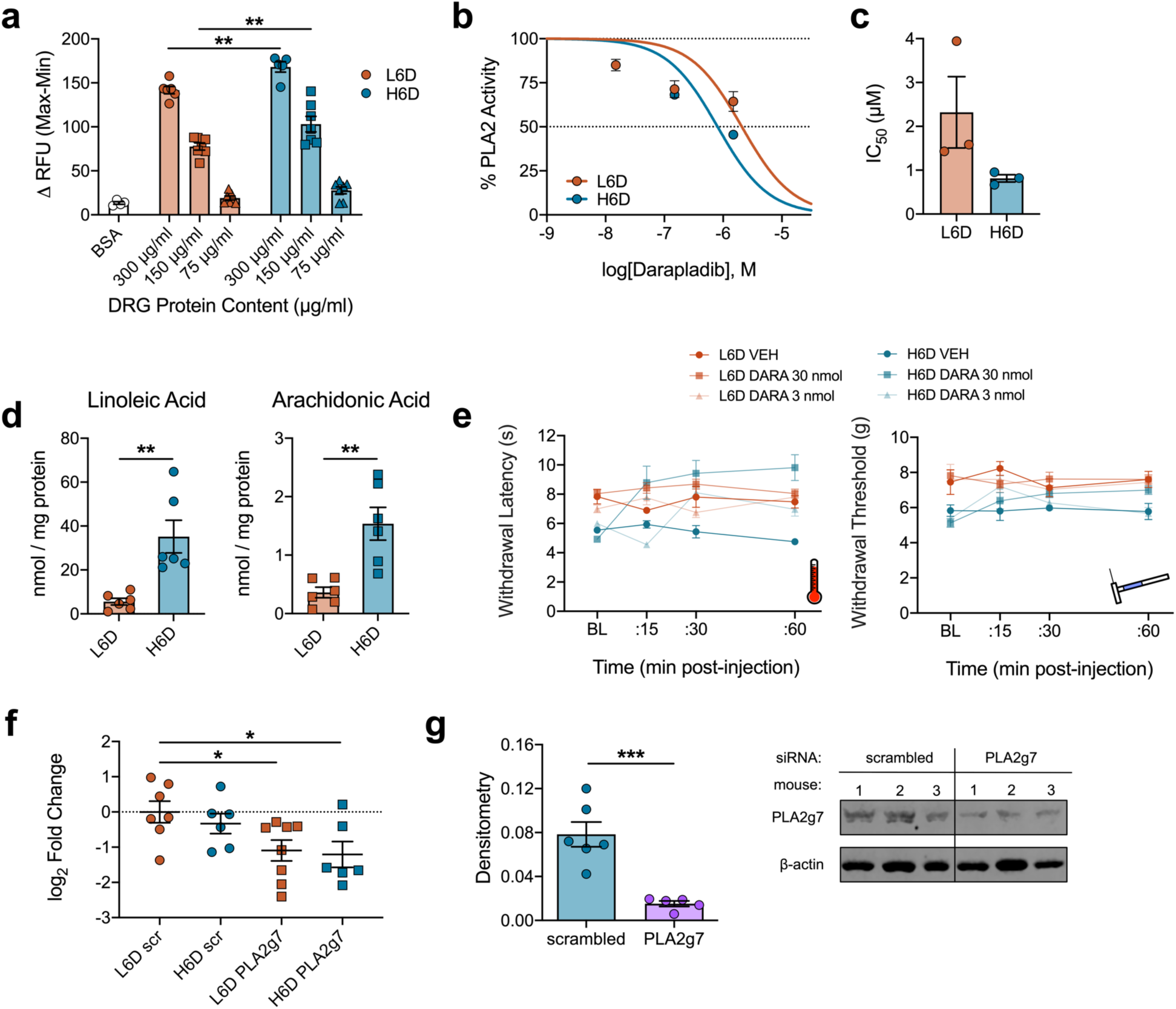
Pharmacological inhibition and genetic knockdown of PLA2g7 in DRG neurons reduces PLA2 activity and attenuates nociceptive hypersensitivities. (**a**) Optimization of the DRG protein concentration used with the PLA2 BODIPY activity assay whereby H6D DRG homogenates demonstrate increased activity at multiple concentrations compared to L6D. **P<0.01 vs L6D, n=5-7 replicates/group. (**b**) Concentration-response curves for darapladib-mediated inhibition of PLA2 activity for H6D and L6D DRG homogenates. (**c**) Half maximal inhibitory concentrations (IC_50_) of darapladib in the PLA2 activity assay as determined by nonlinear regression of the concentration-response curves, n=3 mice/group. (**d**) Total LA and AA levels determined from glabrous hindpaw skin punches for mice on either the H6D or L6D. **P<0.01 vs L6D, n=6 mice/group. (**e**) Time course of dose effects of intraplantar darapladib on heat- and mechanical-evoked nociception. Bold lines represent the group means ± SEM determined from individual animal responses, n=8-9 mice/group. (**f**) qPCR data showing reduced transcript expression of PLA2g7 in lumbar DRG extracts following intrathecal siRNA treatment (q.d. × 3). *P<0.05 vs L6D scrambled, n=6-7 mice/group. (**g**) Western blot analysis of PLA2g7 expression in lumbar DRG of H6D mice following siRNA treatment. ***P<0.001 vs H6D scrambled, n=5-6 mice/group.

**Extended Data Figure 7:**
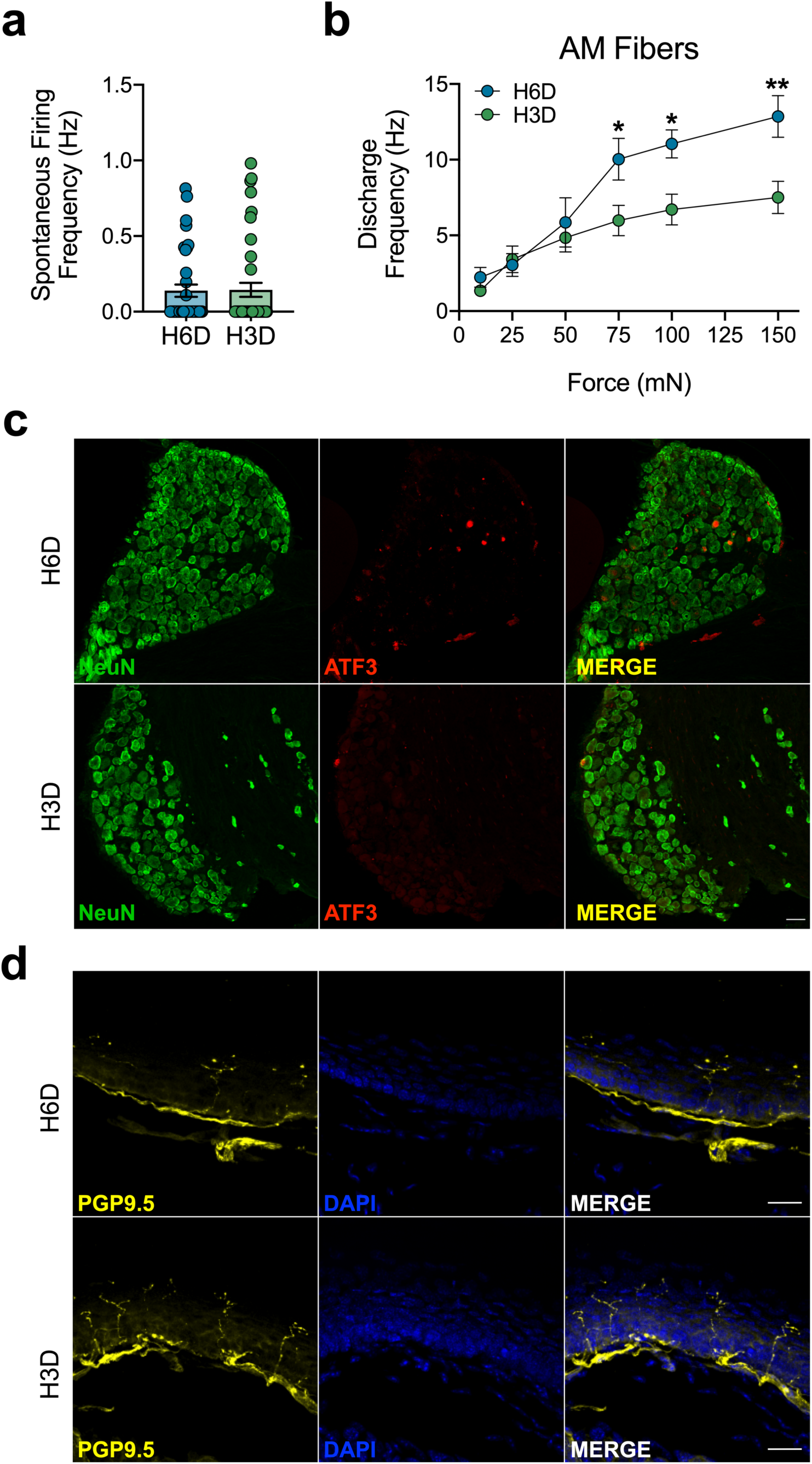
H3D reverses H6D-induced changes in afferent fibers. (**a**) Mean discharge frequencies of spontaneously-active fibers, n=31-40 recordings/group. (**b**) Mean discharge frequencies of AM fibers, n=15-21 fibers/group. (**c**,**d**) Representative immunofluorescence staining of (**c**) ATF3 expression in lumbar DRG neurons and (**d**) glabrous hindpaw skin IENFs in H6D and H3D mice. Scale bars: 50 μm (c) and 25 μm (d).

**Extended Data Figure 8:**
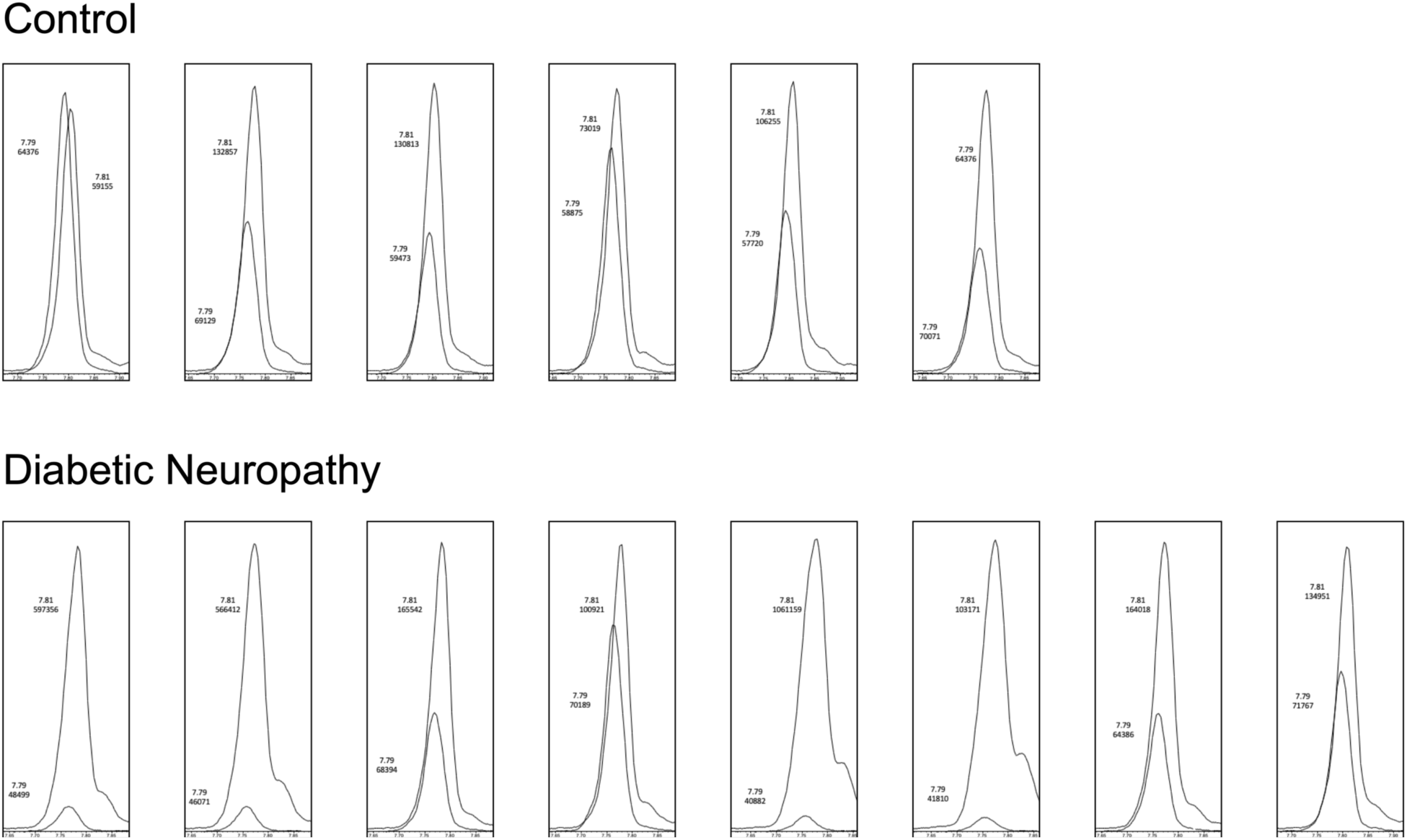
Increased LA content in skin of diabetic neuropathy patients. Chromatogram snapshots of the endogenous LA peak (largest peak, top values) overlaid with the LA-d_4_ internal control peak (smaller peak, bottom values) for skin biopsy extracts from all diabetic neuropathy and control patients. Paired values reflect the retention time (in minutes, top) and integrated area (a.u., bottom) for each peak. Multi-reaction monitoring (MRM) channels in negative ion mode were used for both LA and LA-d_4_, with the collision energy set at 1 eV. Since LA does not produce detectable fragment ions, the channel utilized the parent m/z of 279.2 for the fragment ion as well, thereby excluding species of the same mass that do yield fragment ions.

## Notes

### Competing Interest Statement

The authors have declared no competing interest.

